# Combinatorial Design Testing in Genomes with POLAR-seq

**DOI:** 10.1101/2024.06.06.597521

**Authors:** Klaudia Ciurkot, Xinyu Lu, Anastasiya Malyshava, Livia Soro, Aidan Lees, Thomas E. Gorochowski, Tom Ellis

## Abstract

Synthetic biology projects increasingly use modular DNA assembly or synthetic in vivo recombination to generate diverse combinatorial libraries of genetic constructs for testing. But as these designs expand to multigene systems it becomes challenging to sequence these in a cost-effective way that reveals the genotype to phenotype relationships in the libraries. Here, we introduce a new quick, low-cost method designed for assessing combinational designs of genome-integrated multigene constructs that we call <u>P</u>ool <u>o</u>f <u>L</u>ong <u>A</u>mplified <u>R</u>eads (POLAR) sequencing. POLAR-seq takes genomic DNA isolated from library pools and uses long range PCR to amplify target genomic regions up to 35 kb long containing combinatorial designs. The pool of long amplicons is then directly read by nanopore sequencing with full length reads then used to identify the gene content and structural variation of individual genotypes in the library and read count indicating how abundant a genotype is within the pool. Using yeast cells with loxP-containing synthetic gene clusters that rearrange in vivo in the presence of Cre recombinase, we demonstrate how POLAR-seq can be used to identify global patterns from combinatorial experiments, find the most abundant genotypes in a pool and also be adapted to sequence-verify gene clusters from isolated strains.

## Introduction

Synthetic biology and metabolic engineering aim to create engineered host organisms that perform new tasks or produce a high-value compounds. Typically, this genetic engineering centres around inserting sets of genes and regulatory elements into the genome, usually building-up a cluster of synthetic genes at a single locus^1,2^. For metabolic engineering, constructing multi-step enzymatic pathways requires the balance between achieving sufficient production of the target molecule, achieving cofactor equilibrium, while also minimizing metabolic burden and the accumulation of toxic compounds in the cell^3^. The classic approach to this relies on design-build-test-learn cycles where a cell factory is built and optimized in iterative cycles, an altogether time-consuming task^4,5^. In response to this, many tools have emerged from synthetic biology that utilise modular DNA assembly to rapidly build combinatorial libraries of genetic designs^6,7^. When combined with functional screening assays, these tools enable researchers to test the optimal composition of genes for pathways and genetic circuits, allowing scientists to evaluate a large number of parts in a single experiment^3,4,8,9^. As well as determining the DNA part choices that optimise the transcriptional^3,10–12^ and translational^13,14^ levels of genes (e.g. via promoter and 5’UTR choices), combinatorial libraries also allow researchers to test the impact of the position and orientation of each gene within a cluster or set^15^.

Combinatorial diversity of a set of genes encoding an engineered function is typically achieved during the DNA assembly steps, using pools of parts to generate pools of gene-encoding plasmids^3,8,16^. The final pool of multigene plasmids is then transformed into the host cell, with the DNA ideally being designed so that the cluster of genes ends up stably integrated into the host genome at a known locus for reliable expression. However, an alternative approach for combinatorial library construction is to first place all the genes and regulatory elements for a function into a locus in the host genome and then use targeted recombination at this locus to generate combinatorial diversity *in vivo*, so that the initial synthetic gene set quickly becomes thousands of different designs in the growing cell population.

The widest-known approach for *in vivo* recombination of synthetic genes is SCRaMbLE (Synthetic Chromosome Rearrangement and Modification by LoxPsym-mediated Evolution) where heterologous expression of Cre recombinase rearranges DNA within synthetic gene regions that contain the LoxP sites that it recognises^17^. Originally, SCRaMbLE was developed for the Sc2.0 project, a collaborative project to construct a *Saccharomyces cerevisiae* strain with synthetic chromosomes and ultimately a fully synthetic genome. Following the Sc2.0 design, synthetic chromosomes contain loxPsym sites within the 3’ UTR of all non-essential genes^18^, and these enable inducible Cre-mediated gene deletion, duplication and rearrangements within synthetic chromosomes *in vivo*.

But while the original use of SCRaMbLE was to trigger genome-wide deletions in yeast to better understand gene function, it has also demonstrated use for those seeking to rapidly optimise genetic design of specific functions encoded by synthetic genes and gene clusters. For example, SCRaMbLE in yeast has been used to generate a combinatorial library of strains encoding β-carotene biosynthesis pathway designs, within which strains with a 5-fold increase in β-carotene titres were identified^19^. By having LoxPsym sites flanking the four enzyme-encoding genes, SCRaMbLE shuffles the gene position, orientation, and copy number, leading to diverse designs with altered gene expression^19,20^. More recently, a SCRaMbLE-inspired *in vivo* optimisation system developed by Cautereels *et al*. has been described that enables Gene Expression Modification by LoxPsym-Cre Recombination (GEMbLeR). This system places genes with a panel of regulatory elements into a locus in the yeast genome and uses Cre recombinase to shuffle which promoter and terminators DNA parts flank a gene, diversifying its expression in a population of cells^20^.

With combinatorial library approaches increasingly used in synthetic biology to optimise for engineered novel functions and high yields of biosynthetic products, a critical bottleneck that has emerged is the speed in which the genotypes can be determined. In combinatorial approaches, a phenotypic or fluorescence-associated screen is used to assay thousands of strain designs from a library to isolate the best performing cells^21–23^. The underlying DNA sequence of these cells is then determined to identify the genotype-to-phenotype relationships that explain the best designs. This approach works well when the DNA region undergoing combinatorial design is short and so can be resolved by Sanger sequencing methods (up to 1 kb) or by short-read Illumina amplicon sequencing (up to 500 bp)^24^.

However, when combinatorial DNA constructs introduced into cells are multi-gene length these methods fall short, and this is especially true when the library DNA is integrated into the host genome or is a cluster of genes diversified by SCRaMbLE and related methods. Typically, the researcher is left with no choice but to perform whole-genome sequencing of strains just to resolve the best-performing designs, despite the DNA encoding the synthetic regions being a tiny fraction of the genome. Due to the high cost of this, only a few strains are usually sequenced, meaning that all the sequence-to-phenotype information associated with all other possible designs, including the worst performers, is unknown: representing a major loss for using any downstream learning approaches, including machine learning. Clearly there is a major gap between our abilities to affordably sequence a few engineered genomes in the million bp scale and millions of combinatorial DNA designs that are under 1 kb in length. A targeted method that sequences hundreds or thousands of strains, but only in the region of the genome where combinatorial diversity is introduced is needed.

To help bridge this gap, O’Connell and co-workers recently described a two-step sequencing method where multi-gene length combinatorial library DNA is designed to include a DNA barcode and the plasmid pool post-library construction is subjected to long read sequencing to determine the DNA parts combination associated with each barcode. Following phenotypic screening of the plasmid library in cells, the short barcode region is then amplified for short-read sequencing and the barcode associated with each cell is then mapped to pre-sequenced longer design^25^. While this approach is especially powerful, it requires investment in multiple rounds of sequencing. A more elegant and potentially cheaper approach would be to simply just use long-read sequencing to reveal the full sequence of the DNA in the screened cells without any pre-requirements. This could take advantage of the ability to use barcoding primers with nanopore sequencing so that DNA constructs derived from different cells could be multiplexed in single sequencing run. As an alternative to commercial barcoding kits, Currin et al. proposed amplification of a construct up to 10 kb with tailed primers encoding unique barcodes^26^. Sequencing solutions also exist for combinatorial libraries of plasmids, isolated from individual cell cultures or pools and prepped using transposase-based chemistry^27,28^. The crucial consideration in screening construct libraries is sequencing solely the target region, however methods remain limited for genomically integrated constructs exceeding a standard PCR range.

Here, we introduce a new sequencing method specifically designed to resolve combinational designs of genome-integrated multi-gene DNA constructs that we call <u>P</u>ool <u>o</u>f <u>L</u>ong <u>A</u>mplified <u>R</u>eads (POLAR) sequencing. POLAR-seq, demonstrated here using engineered yeast libraries, relies simply on just three experimental steps; (1) isolation of high molecular weight genomic DNA from cells screened from combinatorial libraries, (2) ultra-long PCR amplification of the engineered genomic region, and (3) long-read nanopore sequencing (**Figure 1A,B**). Reads covering the full length of the engineered region are then selected based on presence of primer binding regions using Porechop^29^ (**Figure 1C**) and reads shorter than the size of the smallest amplicon detected on an agarose gel are removed. Genotypes are revealed by annotating each read with Liftoff^30^, allowing the arrangement and content of the DNA parts in the synthetic region of the genome in the cell population to be analysed with custom scripts.

**Figure 1:**
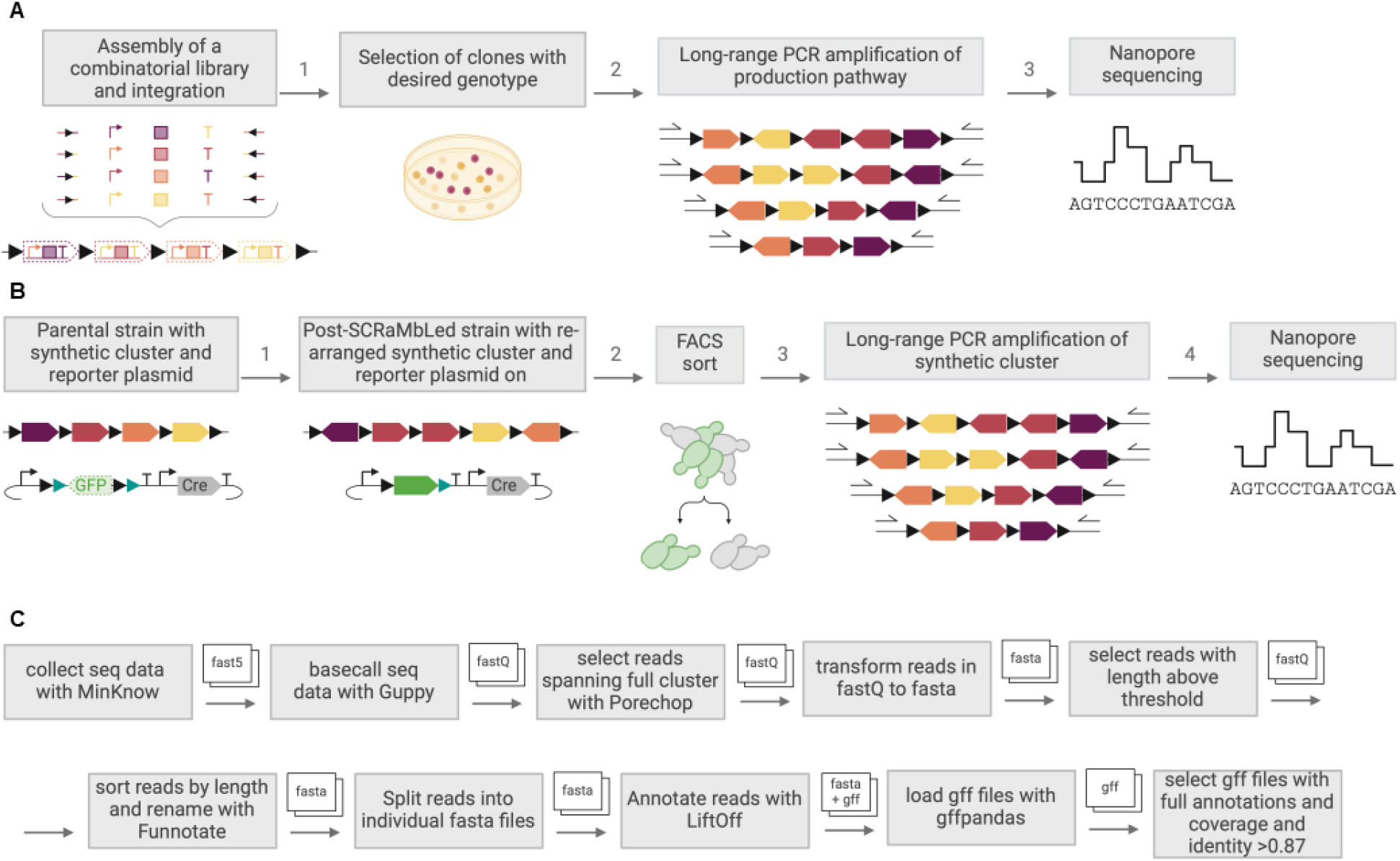
Experimental steps and data analysis of POLAR-seq. **A**. Overview of POLAR-seq workflow for combinatorial libraries: (1) Parts for composing of a gene expression cassette are assembled in a one-pot reaction to create a diverse library of constructs inserted into the genome. (2) Colonies of desired phenotype are selected for screening. (3) Cells are grown and subjected to genomic DNA isolation which is used as a template in long-range PCR to amplify synthetic cluster. (4) PCR amplicons are prepped for sequencing using the ligation kit and sequenced on nanopore platform. **B**. Overview of POLAR-seq workflow for *in vivo* recombined libraries: (1) Parental strain with a synthetic cluster that contains a set of genes (colour) flanked by loxP sites (black triangle) is subjected to SCRaMbLE. Cells also contain a SCRaMbLE reporter plasmid encoding Cre recombinase (in grey) and flipped GFP reporter (green) between two pairs of loxP sites (black and green triangles). Cre-mediated recombination leads to rearrangements within the cluster and reversal of GFP orientation (green) which turns on cell fluorescence. (2) GFP positive cells that underwent SCRaMbLE re-arrangements can be retrieved by FACS. Steps (3) and (4) are performed as described in panel A. **C**. Workflow for data analysis steps post-sequencing and the resulting files, as described in the methods. Figure created with BioRender.com.

## Results

### Establishing long range PCR from engineering yeast genome samples

To develop a method to selectively sequence DNA from a defined region of a host genome, we focused on optimisation of conditions for long range PCR from genomic targets, specifically testing this with a *S. cerevisiae* yeast strain containing a 28 kb synthetic gene cluster integrated into the *URA3* locus at chromosome V. Long range PCR amplification is possible up to 30 kb and beyond, but is known to be challenging for several reasons, especially if the DNA used as a template has poor integrity. Therefore, to prevent the use of damaged DNA and to remove any PCR inhibitors from the template DNA substrate, our first priority was to employ a genomic DNA isolation method that maintains high-molecular DNA fragments (>50 kb), as described in the methods section. The method of Denis et al. allows to extract yeast DNA with N50 of 50 kb while maintaining purity suitable for long reads sequencing. ^31^

Another known challenge for long range PCR is the presence of secondary structure and high GC content in the DNA to be amplified, as both can prevent primers from binding correctly and can cause premature termination of strand elongation. GC content was not expected to be of concern for the work described here, as the clusters we amplified had a normal 40% GC content. A further challenge is the design of primer pairs specific for the desired target, where it is important that designs do not generate non-specific amplicons. To avoid this, we therefore initially examined three sets of primers with different binding regions.

Finally, selection of a DNA polymerase is crucial for successful generation of ultra-long PCR amplicons (**Figure 2A**)^32^. For this purpose, we tested a range of commercially available DNA polymerases expected to be able to do long amplicons. Two of the tested polymerases, LA Taq HS and Herculase II generated PCR products from 18 kb up to 35 kb (**Figure 2B**). LA Taq HS is a mix of Taq polymerase and a DNA polymerase exhibiting 3’→5’ exonuclease activity with hot-start mediated by a monoclonal antibody against *Taq* Polymerase. In the further experiments we continued all work with LA Taq HS which achieved higher PCR product yields compared to Herculase II (**Figure 2B**). Importantly, all PCRs were also set to a total volume of 25 µL to avoid any thermal gradients. PCRs were then further optimized to find conditions that prevent formation of unspecific amplicons, and this led to us a protocol with primer concentrations reduced to a final concentration of 0.1 µM and the amount of template DNA provided to each reaction reduced to 20 ng (**Figure 2C, Supplementary Method 2**).

**Figure 2:**
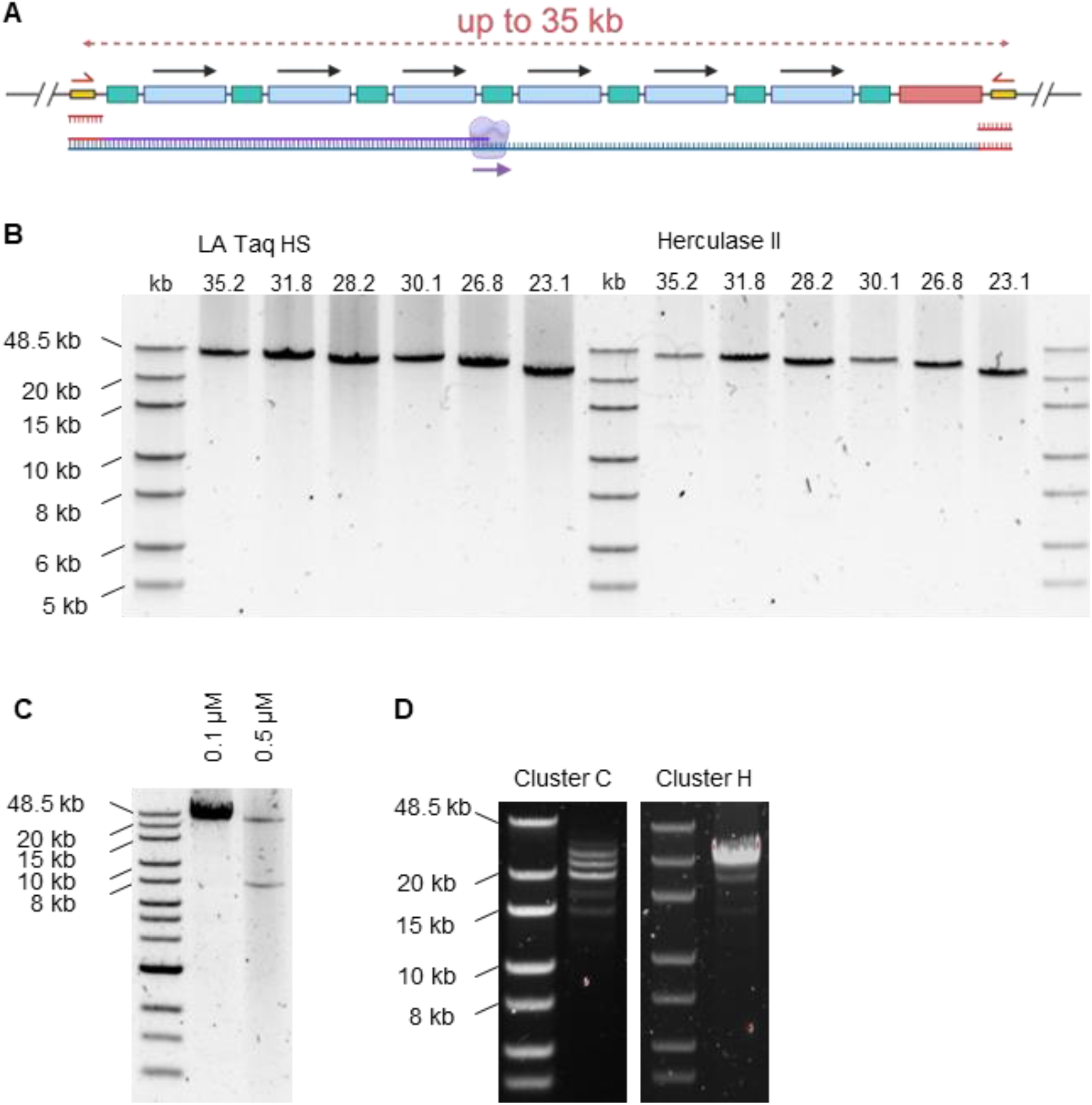
Long-range PCR amplification of genomic regions. **A**. A multigene synthetic construct integrated into a genomic locus with a selectable marker gene (red) is flanked by primer binding sites (yellow) that are used for PCR amplification of clusters up to 35 kb in length. **B**. Comparison of LA Taq Hot Start and Herculase II polymerases for amplification of 23 to 35 kb regions using primers KC001/2, KC003/4, KC005/6. The annealing temperature of all PCR reaction with Herculase II was set to 52.5 °C. Extension time was set to 15 min for all PCR products generated with LA Taq, while this parameter was adjusted for reactions with Herculase II (30.1 – 35.2 kb: 18 min, 23.1 – 28.2 kb: 14 min). Amplicons were diluted x10 and 10 µL were loaded on the gel. PCR amplification with KOD Xtreme produced unspecific products and repliQa HiFi Tough failed to generate any products (not shown). **C**. Optimization of PCR amplification with LA Taq HS. Reduction of primers concentration from 0.5 µM to 0.1 µM prevents formation of unspecific products. **D**. PCR amplification of C and H clusters from pools of cells subjected SCRaMbLE and FACS-sort.

### Long range PCR from libraries of yeast cells with rearranged genomic DNA

Having established a protocol for long-range PCR from isolated yeast genomic DNA, we next set out to show that this can amplify long-range DNA from synthetic gene clusters in the yeast genome when taking diverse pools of yeast cells, rather than just taking from single colonies of yeast cultured to high density. To demonstrate this we grew two yeast strains, known as strain C and strain H, which have each been constructed for other projects to have a synthetic multi-gene cluster integrated into the *URA3* locus of their genomes (see **SI Note 1**). Strain C’s cluster is 28 kb long and contains 9 genes with roles in the control of the cell cycle in yeast, an essential function but with gene redundancy. Strain H contains an 18 kb cluster of 7 genes encoding a biosynthesis pathway that is essential to be fully intact in cells grown in minimal media. In both cases the synthetic clusters have been designed with loxP sites between each gene so that genes in the cluster can be rearranged, deleted and duplicated *in vivo* through expression of Cre recombinase in the nucleus. The cluster in strain H uses symmetrical loxP sites (loxPsyms) to allow all types of rearrangements, but the cluster in strain C uses non-symmetrical loxP sites and so heavily favours deletion events.

Cells of strain C and strain H were transformed with a plasmid construct that expresses Cre recombinase and shuttles this into the nucleus to recombine genomic loxP sites when induced by β-estradiol^18^. This plasmid, when used in a SCRaMbLE with Sc2.0 yeast strains, allows for rapid induction of Cre and leads to deletions, duplications, and/or translocations of genes flanked by loxP sites. Strain C and H cells were grown in flasks in minimal media and given β-estradiol to induce SCRaMbLE with the intention of rapidly diversifying the content and arrangement of their cluster genes in the large population of cells in the flask.

Using a SCRaMbLE reporter construct that reverses GFP-encoding ORF DNA into an expressing orientation in response to Cre-Lox recombination, we next sorted the yeast by fluorescence to capture the cells in the population most likely to have had SCRaMbLE events in their genome. Sorting of the GFP+ cells was performed by FACS to isolate population with DNA rearrangements and high-quality genomic DNA from the pools of sorted yeast cells was then obtained. Within these genomic DNA samples, should be the DNA sections encoding the clusters after their diverse rearrangements.

Long range PCR was then attempted as before, but now using pool genomic DNA samples as template. For both the C and H strain pools, the PCR protocols proved successful in amplifying the cluster DNA regions. The PCR of the cluster DNA from the sorted C strain pool led to amplification of DNA fragments of 7 distinguishable size group from 12 kb up to 28 kb in length, the original length of the C cluster. (**Figure 2D**). Each size group represents clusters with a different number of deleted genes. Meanwhile, PCR of cluster DNA from the sorted H strain pool gave amplicon products mostly around 20 kb in length, but with two groups of shorter amplicons (approximately 12 kb and 18 kb) also visible by agarose gel electrophoresis (**Figure 2D**).

### Long-read sequencing reveals genotypes of SCRaMbLEd clusters

We next assessed whether PCR libraries generated from DNA extracted from pools of SCRaMbLEd cells were suitable for nanopore sequencing. Amplicons from long-range PCR from the C strain pool were prepared for sequencing using the ligation kit SQK-LSK109 and sequencing was performed on R9.4.1 Flongle flow cells which allows a rapid, low-cost experiment while providing up to 2 Gb of data. To determine whether sequencing on Flongle underrepresents certain genotypes, we also conducted sequencing of the amplified C clusters on R10.4.1 MinION flow cell.

Sequencing of the pool of C clusters on both types of flow cells was successful and after a series of data analysis steps (see **Methods**) we ended up obtaining over 2000 high-quality annotated reads that cover the full length of the synthetic cluster via the Flongle sequencing, and over 100 times more reads (>245,000) via the MinION sequencing (**Table 1**). In both cases the reads were suitable for determining the C cluster genotypes of the cells from the sorted post-SCRaMbLE pool. Importantly, the sequencing of the long amplicons allowed for rapid identification of Cre/loxP mediated gene deletion combinations within the C cluster. It led to the discovery of 432 unique C cluster genotypes using the MinION flow cell data and 81 unique genotypes using the Flongle data. Importantly, the 81 genotypes recognized from data obtained with Flongle device overlapped with those found via MinION flow cell sequencing, where the least frequent genotypes represented only 0.02% of the total reads and therefore only a tiny fraction of the yeast cells in the sorted pool.

**Table 1:** Overview of the nanopore sequencing data generated and processed for sorted strain pools of SCRaMbLEd C and H clusters.

| Library | C cluster | C cluster | H cluster |
| --- | --- | --- | --- |
| Flow cell | Flongle | MinION | Flongle |
| Chemistry | v9 | v14 | v9 |
| Bases generated [Mb] | 232.06 | 26,850 | 658.75 |
| Reads passed | 11,765 | 1,820,119 | 36,986 |
| Reads demultiplexed | 3,341 | 413,702 | 6,635 |
| Reads above threshold | 2,839 | 332,090 | 6,401 |
| Reads annotated | 2,084 | 245,895 | 4,050 |
| Detected genotypes | 81 | 432 | 127 |

Next, the set of 10 most abundant genotypes in the data were visualized and the count of reads corresponding to each genotype was compared between runs (**Figure 3A**). These 10 genotypes together account for around 75% of all reads from the sequencing runs and all contain at least one gene deletion from the C cluster, with some showing up to 4 deleted genes. For all 10 genotypes the percentages of reads from the MinION and Flongle experiments were in close agreement, with variability between sequencing runs only expected to be caused by pipetting errors, for example while setting up the PCRs and loading the sequencing flow cells. This shows that either flow cell is suitable for these experiments. Genotypes *i*–*iii* were by far the most abundant, together accounting for 45.8% of MinION reads and 47.2% Flongle reads, presumably due to the cells with these genotypes being most abundant in the sorted cell pool. Overall, this initial analysis shows that the frequency of SCRaMbLE rearrangements in a synthetic yeast cluster can be estimated from pooled samples by long-range PCR and long read sequencing.

**Figure 3:**
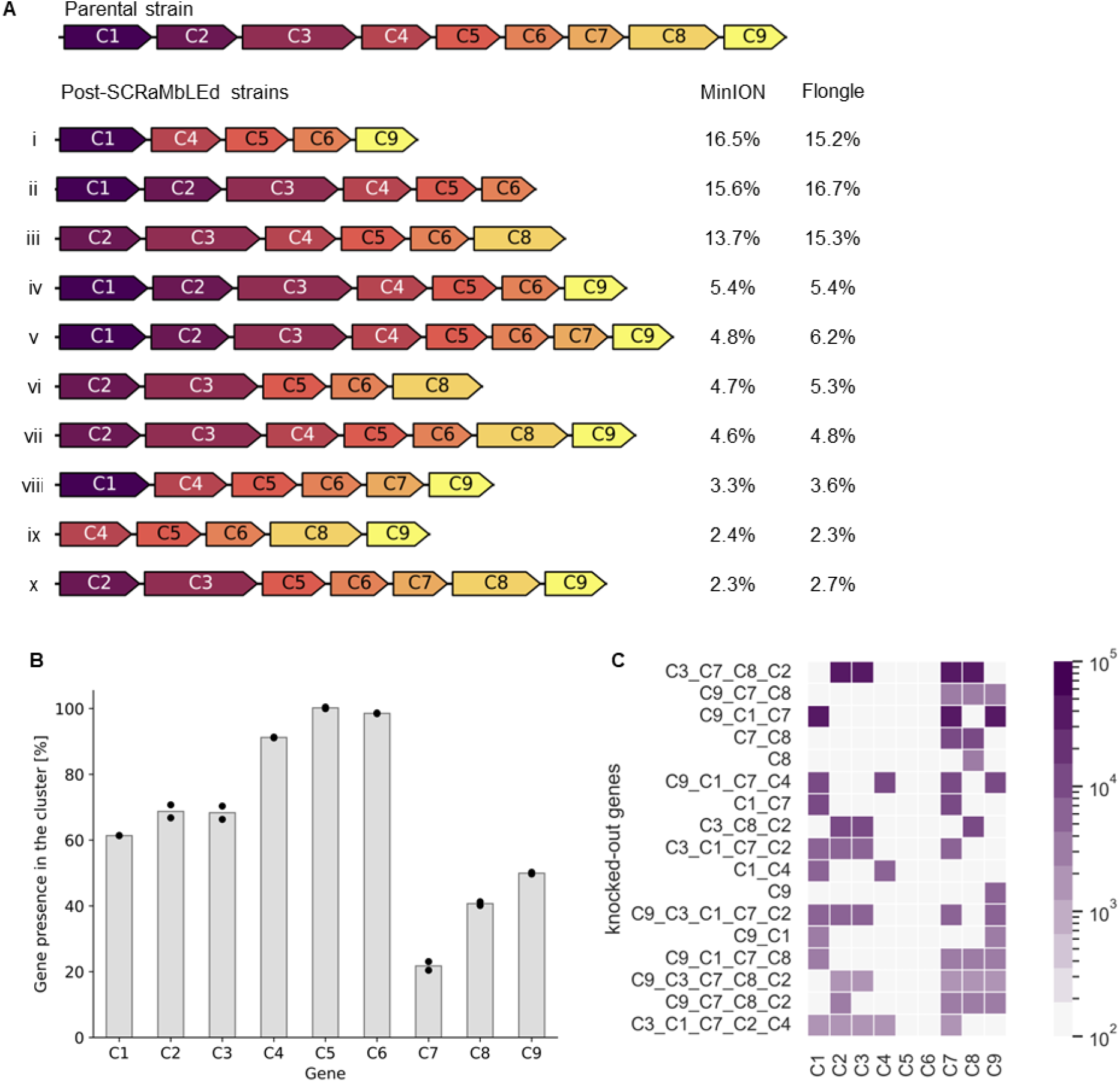
POLAR-seq of the C cluster subjected to combinatorial deletions. **A**. Genotype of the parental strain and ten most abundant rearranged genotypes discovered by sequencing on MinION and Flongle flow cells. The percentage represents number of reads discovered for a given genotype and normalized by all the annotated reads. **B**. Frequency of genes maintained in the C cluster after combinatorial deletions expressed as percentage (count of gene annotation normalized by the total number of annotated reads). Data points represent frequency from the two independent sequencing runs (MinION and Flongle). **C**. Combinations of gene knockouts recorded in rearranged C clusters. Heat map was prepared for the 20 most abundant genotypes. The colour intensity indicates the number of reads detected for the given genotype.

### Rearrangements within the cluster can be tracked globally

Over 99.5% of the reads from the C cluster pool contained a gene deletion, with only 4.4% additionally containing a gene duplication. To investigate which genes are deleted from the cluster most frequently, we counted the occurrence of each gene in all annotated reads (**Figure 3B**). The frequency of genes in the cluster obtained from the MinION and Flongle sequencing closely matched. As shown in the **Figure 3B**, the most frequently deleted gene was C7 (found in only 20% of the detected genotypes), followed by C8 (40%) and C9 (50%). Whereas genes C4, C5 and C6 were those most frequently kept in the cluster; with C4 and C6 gene detected in 91% and 98% of the reads, respectively, and gene C5 being found in 100% of the reads. This kind of analysis is especially useful for quantifying gene essentiality, for example in a growth condition. Gene C5 in this case proving to be essential (found in 100% of reads), and gene C7 proving to be non-essential (only found in 20% of reads).

Importantly, long-read sequencing also reveals the combinations of genes that can be deleted within the same cluster, something difficult to achieve from a pooled sample by short-read sequencing or by RNAseq. Analysis of common gene deletion combinations (**Figure 3C**) showed that genes C7 and C1 were commonly deleted together, as were C7 and C8, and also genes C2 and C3. Visualization of deletion combinations also reveals those rarely or never deleted together, for example genes C4 and C8.

### POLAR-seq allows to study gene inversions and translocations

Having shown with the C strain that POLAR-seq identifies deletion genotypes in a post-SCRaMbLE library, we tested whether it serves to identify other structural variations. For this we used the H strain, where the cluster genes are flanked by symmetrical loxPsym sites that permit gene duplication, translocations and inversions as well as deletions. Amplified cluster DNA from a post-SCRaMbLE library of these cells was sequenced on Flongle and data analysis from this gave over 4000 cluster-length annotated reads that identified 127 different genotypes (**Table 1**). These were then classed based on the type of rearrangements seen in the reads (**Figure 4A**) with the most common being genotypes with deletions, duplications and inversions all occurring (48% of all genotypes) and the second most common being genotypes with just duplications (39%). Deletions were almost never seen because all 7 genes in the cluster are essential for cell viability in the conditions they were cultured in.

**Figure 4:**
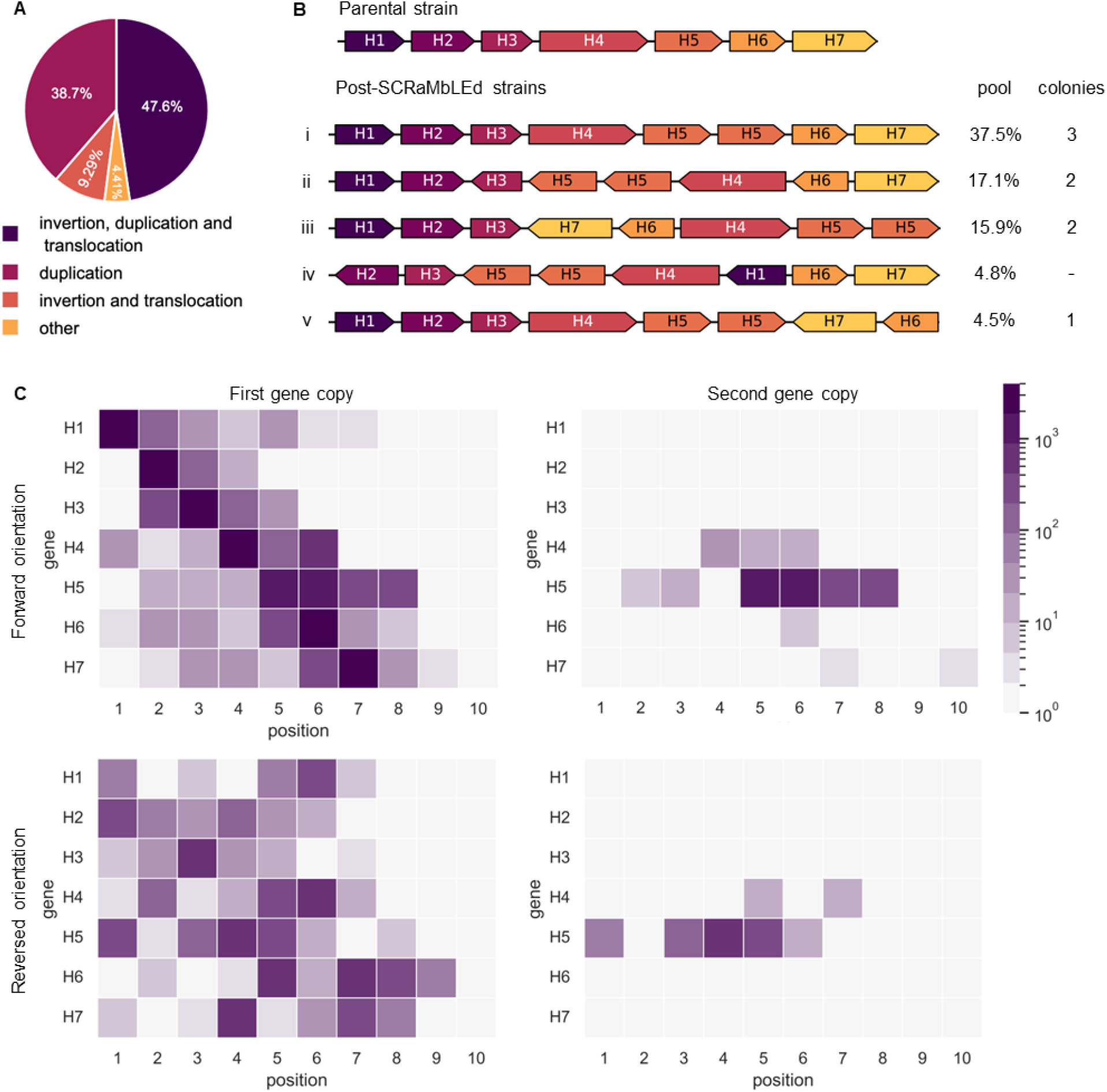
POLAR-seq of the H cluster subjected to SCRaMbLE. **A**. Types of detected re-arrangements in the H cluster. Each genotype was classified depending on re-arrangement type and the number of reads representing each type was normalized by the total number of annotated reads. **B**. Genotype of the parental cluster and top five post-SCRaMbLE genotypes detected from a pool of FACS-sorted cells, either normalized by the total number of annotated reads (*pool*) or from 8 randomly picked colonies from the pool (*colonies*). **C**. Relationship between gene copy number, gene orientation, and gene position in clusters sequenced from the post-SCRaMbLE pool of cells. Each heatmap shows the frequency of finding each gene (H1-H7) in positions 1 through 10 of a sequenced cluster, where the parental strain has genes H1-H7 in positions 1 to 7 respectively, all in forward orientation and single gene copy. Left heatmaps show the frequency of finding the first/only copy of a gene in each position in a forward (top) or reverse (bottom) orientation. Right heatmaps show the frequency of finding genes in each position that are duplicated in forward (top) or reverse (bottom) orientations.

In total 92% of all reads showed a duplication of gene H5, and this was reflected in the 5 most abundant genotypes found in read data (**Figure 4B**). Genotype *i* showed just a single duplication of H5 while the other four top genotypes (*ii-v*) showed duplication in association with inversion and/or translocations of the other genes Further analysis based on all of the determined genotypes was used to reveal the frequency of each gene being found to have moved position in the cluster, inverted its orientation or being duplicated (**Figure 4C**). This kind of global analysis of gene rearrangements has not been demonstrated before in any published SCRaMbLE experiments and is not possible to do by standard PCRTag analysis^33^ or the recently described LoxPTag^34^ method. POLAR-seq in particular shows its advantage in detecting duplications, which are not easily identifiable by non-sequencing approaches. Importantly, the high frequency of detecting duplications in the dataset here also relieves concerns that the PCR-based target enrichment used by POLAR-seq might struggle with clusters with repetitive DNA.

Finally, to confirm whether genotype abundance is consistent with the single cells, we next sequenced 8 randomly picked yeast colonies plated out from the same post-SCRaMbLE library. Colonies were cultured in 2 mL of YPD medium, genomic DNA isolated and cluster sequence PCR amplified using LA Taq polymerase and primers KC007 and KC015. These primers match the primers KC005 and KC006 used in POLAR-seq but have an additional 25 base tail (**SI Table 1**) that serves as a barcode for demultiplexing sequencing reads from combined samples, according to the method published by Currin et al^35^. Three of the tested colonies exhibited genotype with a H5 duplication (*i*), which was found in 37.5% of reads generated from sequencing pool (**Figure 4B**). The next most abundant two genotypes (*ii, iii*), were detected in 2 colonies each and genotype *v* was found in 1 colony. Genotype *iv* was not found among 8 sequenced colonies. Overall, the frequency at which cluster genotypes were present in this small random selection of colonies broadly matched the abundance of reads for the genotypes from our POLAR-seq experiment, giving us confidence the read numbers in our analysis are representative of the number of cells with that genotype in the sampled pool.

## Discussion

Here, we present a method for long-range amplification and long-read sequencing of genome-integrated synthetic DNA regions that is suitable for a pool of FACS-sorted cells. By sequencing the DNA of an entire pool of cells, a larger population can be screened to identify rare genotypes, which may not be detected by analysis of individually isolated strains. When applied to SCRaMbLEd synthetic gene clusters in yeast, the POLAR-seq method allows rapid and efficient identification of structural rearrangements within the cluster including duplications, deletions, inversions, and translocations. As the method simply requires just genomic extraction, PCR amplification and then low coverage nanopore sequencing, it is quick and affordable. Considering the cost of reagents, kits and flow cells, the method reveals yeast cluster genotypes from a sorted library at an approximate cost of $0.09 per genotype when using a MinION flow cell, and $1.5 per genotype when using a Flongle, assuming that 25 annotated full-length reads is sufficient to identify a cluster. A Flongle POLAR-seq experiment costs only $150 in total, and can also be adapted as in Figure 4b to sequence barcoded amplified DNA from up to 24 isolated colonies, representing a cost of less than $10 per strain to determine individual cluster genotypes.

In developing this method, we demonstrated the use of POLAR-seq to study Cre-mediated rearrangements and deletions within loxP site-containing synthetic clusters in *S. cerevisiae* cells showing that the method was especially suited for revealing duplications, a rearrangement difficult to assess with classic short read sequencing approaches. POLAR-seq is however, not limited to just structural rearrangement applications. POLAR-seq could also be used to reveal the optimal part composition of any DNA encoded constructs so long as the desired function of this can be screened and retrieved, e.g., by FACS-based cell sorting. For example, instead of rationally selecting of promoter parts for each gene in a multigene metabolic pathway, all DNA parts for this pathway could be assembled in a one-pot reaction, integrated into yeast and the optimal designs for function can then be determined by combining a functional screen with POLAR-seq. A further possibility afforded by POLAR-seq is in quantifying the relative fitness of each genotype within a library. Genotypes with the highest percentage of reads correspond to the strains with the highest abundance in the sampled pool (**Figure 4B**), and so performing POLAR-seq at sequential time points when a library pool is grown in a condition of interest could be used to reveal the strain genotypes that outgrow others due to better fitness. Being able to measure function and fitness from thousands of variants in a single experiment is likely to open up new opportunities in machine learning for synthetic DNA design.

A limitation of PCR amplification, however, is frequency of introduced errors, especially for low fidelity polymerases such as Taq. Therefore, we would recommend against using POLAR-seq to determine variation in libraries at the single base level, rather than at the parts and structure level as used here. Base mutations do not prevent our analysis from correct recognition of cluster’s genes as this has been set to accept up to 12.6% of erroneous bases within a sequence. In applications sensitive to SNPs, the PCR step in POLAR-seq can be improved using the high fidelity Herculase II polymerase (1 error in 777 kb). Another method to improve quality of sequencing data is the chemistry of the selected library prep kit and the flow cell. During the preparation of this manuscript, only v9 Flongle flow cells were available, however these can be now substituted with v14 flow cells. With this advance, the raw read accuracy in simplex now is reported to exceed Q28 (99.8%)^36,37^.

Finally, the POLAR-seq method is limited by the size of the region of interest, as this must be suitable for PCR amplification. As it stands POLAR-seq is not suitable to study content and structural diversity in genomic regions with lengths beyond 35 kb. Other methods available for region-specific enrichment of genome sequencing include Oxford Nanopore Technology’s adaptive sampling method, and CRISPR/Cas9-based ‘catching’ of specific genomic regions^38–40^. However, these work by exclusion of unwanted sequence rather than amplification of target DNA, meaning that orders of magnitude more genomic DNA needs to be harvested from samples to achieve the same coverage. PCR amplification is therefore much better suited to this task, and hopefully with continued improvements in long range DNA polymerases, or by adapting rolling-circle amplification methods, the lengths of genomic regions suitable for POLAR-seq analysis will increase in the future.

## Acknowledgments

This research was supported by a Wellcome Trust Discretionary Award (221267/Z/20/Z) providing funding for K.C. and T.E., a Chinese Scholarship Council (CSC) PhD scholarship to X.L., and a Darwin Trust of Edinburgh PhD scholarship to A.M. T.E.G. was supported by a Royal Society University Research Fellowship grant URF\R\221008 and a Turing Fellowship from The Alan Turing Institute under EPSRC grant EP/N510129/1.

## Methods

### Strains and Media

All yeast strains used in this study are derivatives of BY4741 yeast (*MATa his3Δ1 leu2Δ0 met15Δ0 ura3Δ0*). Yeast strain C is a haploid strain generated by genetic engineering for an independent research project and has a synthetic cluster of 9 cell cycle related genes assembled and integrated into the *URA3* locus on chromosome V. Yeast strain H is also a haploid strain with a synthetic cluster of 7 metabolism related genes assembled and integrated into the *URA3* locus.

Yeast extract Peptone Dextrose (YPD) media (10 g L^-1^ yeast extract (VWR), 20 g L^-1^ peptone (VWR), 20 g L^-1^ glucose (VWR)) was used for culturing of yeast strains without the SCRaMbLE reporter, unless otherwise stated. Synthetic Complete media (SC; 6.7 g L^-1^ Yeast Nitrogen Base without amino acids (Sigma Aldrich), 1.4 g L^-1^ Yeast Synthetic Drop-out Medium Supplements without L-uracil, L-tryptophan, L-histidine, L-leucine, 20 g L^-1^ glucose (Sigma Aldrich)) was used for auxotrophic selection. Amino acids such as 20 mg L^-1^ L-tryptophan, 20 mg L^-1^ L-histidine, 20 mg L^-1^ uracil and 120 mg L^-1^ L-leucine were supplemented into SC media depending on the required auxotrophic selection. For growth on plates, media were supplemented with 20 g L^-1^ bacto-agar (VWR).

### SCRaMbLE

C and H yeast strains were transformed with plasmids for β-estradiol-induced Cre expression and for GFP expression in response to nuclear Cre activity. Strains were then grown overnight in 2 mL SC complete media with appropriate selection (30°C, 250 rpm). Cultures were diluted to an OD_600_ of ∼0.2 in 5 mL SC complete media with appropriate selection and grown for 4 hours. SCRaMbLE was induced by addition of β-estradiol (dissolved in DMSO) to a final concentration of 1 μM. Cultures were grown for an additional 4 hours before being washed twice in water and resuspend in 1 mL PBS for Fluorescence-Activated Cell Sorting (FACS).

### Fluorescence-Activated Cell Sorting (FACS)

FACS sorting was performed on the BD FACSAria III Cell Sorter (BD Biosciences) to select for yeast cells with GFP fluorescence. Yeast cells, washed and resuspended in PBS buffer post-SCRaMbLE, were transferred to a 5 mL FACS tube (Invitrogen) and diluted to appropriate density with PBS buffer for FACS sorting. The 70 μm nozzle was selected for the sorting. Around 1 million GFP+ cells sorted from the FACS instrument were collected in a 15 mL Falcon centrifuge tube. A subset of cells was immediately inoculated into appropriate media and grown for 2-3 days to reach saturation (30°C, 250 rpm). The remaining cells were spun down at 4000 rpm for 20 min. The pellet was resuspended in 0.25 mL PBS and stocked in 25% glycerol (final concentration) at -80°C. For the FACS analysis, yeast cells were firstly selected based on morphology (FSC-A vs SSC-A). Single cells were then selected based on a double doublet-discrimination (FSC-A vs FSC-H and SSC-W vs SSC-H). Single GFP+ cells were then selected based on GFP expression (FSC-A vs GFP). GFP expression was selected with the 530/30 bandpass filter.

### Genomic DNA isolation from yeast

Genomic DNA was isolated according to the method published by Denis et al. ^31^ with following modifications: yeast culture was grown until cells reached OD=5-10, zymolyase was replaced with lyticase (Sigma Aldrich, 600 U per 1 mL of OD=1) and all centrifugation steps were performed at 4000 g (**Supplementary Method 1**). The adapted protocol was also tested in minimized scale, making use of 2 mL of yeast culture and proportionally reduced reagents.

### PCR amplification

Initial testing used four DNA polymerases: LA Taq Hot Start (TaKaRa, Shiga Japan), Herculase II (Agilent, Santa Clara, United States), KOD Xtreme (Sigma Aldrich, Saint Louis, United States) and repliQa HiFi ToughMix (QuantaBio, Beverly, United States). PCR reactions were set up in 25 µL volume according to manufacturers’ protocol using 20 ng of genomic DNA as template and reduced concentration of primers (final concentration 0.1 µM). Amplification was verified by gel electrophoresis on 0.5% agarose gel run at 50 V for 5 h. PCR products were purified using AMPure beads (1.8x volume of the PCR reaction) and quantified using Qubit dsDNA Broad Range kit and Qubit 2.0 Fluorometer. Prior to sequencing, quality of DNA was evaluated by measuring absorbance using Nanodrop spectroscopy.

### Nanopore Sequencing

Pools of amplicons were prepared for sequencing using the NEBNext Companion Module and the ligation kit SQK-LSK109 or SQK-LSK114. The DNA library was sequenced on the Flongle (FLG001) or MinION flow cell (FLO-MIN114) using the MinION Mk1B device. In each case data was collected during sequencing with the latest version of MinKnow (22.05.5 – 23.04.6).

### Data Analysis

#### Basecalling sequencing data and Demultiplexing

Raw data was basecalled with Guppy (6.1.5 - 6.5.7) using high-accuracy model and only reads with Q-score above 9 were kept for further analysis. To select reads that span full amplicons we used the demultiplexing function of Porechop^29^. Reads were demultiplexed to firstly find binding regions of primer KC005 and subsequently of primer KC006 using default settings (barcode_threshold 75 and barcode_diff 5). This step ensures that partial PCR products and reads corresponding to shredded DNA fragments are not further considered.

#### Filtering sequencing reads

Demultiplexed reads in fastq format were next transformed into fasta sequence and filtered based on size. The threshold is set based on the smallest detectable PCR product, specifically 11 kb for the C cluster and 10 kb for the H cluster. The remaining reads were sorted by length and renamed numerically with funannotate^41^. Reads were split into individual fasta files using splitfasta.

#### Annotating sequencing reads

Reads were annotated by Liftoff^30^ from the provided reference sequence that contains all genes present in the parental cluster design. Liftoff generated a gff file for each read and these were uploaded as Python dataframe with gffpandas. This allowed retrieval of the position of each gene and could be also used to determine the orientation of a read based on the fact that both C and H clusters contain an auxotrophic marker at the 3’ end. Thus, the read orientation can be defined by presence of this marker on the forward or the reverse stand. If the marker is on the reverse strand and appears as the last gene, the reverse complement sequence is generated for the given fasta file with BioPython SeqIO library. Subsequently, the generated sequences were re-annotated.

#### Filtering GFF files

The GFF files were then combined for the reads encoding the forward strand and filtered based on the following criteria: 1) the first annotation must be within the defined distance *A* from the start of the read; 2) the gap between two annotations must be lower than distance *B*; 3) the sequence identity and coverage of each gene annotated in a read must be above the 87.4% threshold defined based on Q-score. 4) The reads must encode DNA sequence from the auxotrophic marker as the last annotated gene. For criteria 1 to 3, values were determined based on the expected error of 12.6% when Q-score=9 is set as a threshold for reads binned as passed in high-accuracy base-calling. This corresponds to a threshold of 87.4% of correct bases in each read. Distance *A* (criterium 1) was calculated based on the distance from the start of the amplicon to the first annotated gene plus 12.6% buffer on each (corresponding to the allowed sequencing error when Q-score=9). Distance *B* was calculated as the distance between genes in the cluster plus 12.6% buffer on each end.

#### Data visualization

After filtering, GFF files that fulfilled the abovementioned criteria were loaded as a Python dataframe with gffpandas. Only gene names were maintained and were next merged into a single line, allowing to count the same genotypes. Selected genotypes were visualized with DnaFeaturesViewer^42^. Plots were then created with Python packages: matplotlib, and plotly.

