## Supplementary Information for "Combinatorial Design Testing in Genomes with POLAR-seq"

### TABLE OF CONTENTS

**Supplementary Methods**

Supplementary Method 1: Genomic DNA isolation 2

Supplementary Method 2: Long range PCR amplification 3

**Supplementary Notes**

Supplementary Note 1: Synthetic clusters used in this study 4

**Supplementary Tables**

Supplementary Table 1: List of primers used in this study 5

Supplementary Table 2: Accession numbers of raw sequencing data submitted to SRA 6

Supplementary Table 3: Median Q-scores of the data generated from described sequencing runs. 7

**Supplementary Figures**

Supplementary Figure 1: Position of annotations in the analysed reads from Flongle sequencing 8

Supplementary Figure 2: Combination of the top 11 gene knockouts in the C cluster sequenced on Flongle flow cell. 9

### Supplementary Method 1: Genomic DNA isolation

Adapted methods for genomic DNA isolation are available at:

https://benchling.com/s/prt-TZEDV9nnYDTw808JZ8UG?m=slm-6wfDTBwcVjBCil8Kfx8y

https://benchling.com/s/prt-z5JaWpuZcaV7jrnvNKrj?m=slm-n3miNqSfu9ToHCerEUiC

### Supplementary Method 2: Long range PCR amplification

PCR reaction with LA Taq

| Reaction mix | |
|---|---|
| TaKaRa LA Taq HS pol (5 U/µl) | 0.25 |
| 10x LA PCR Buffer Ⅱ ($Mg^{2+}$ plus) | 2.5 |
| dNTPs (2.5 mM each) | 4 |
| FW primer (2 µM) | 1.25 |
| RV primer (2 µM) | 1.25 |
| Template (20 ng gDNA) | 1 |
| Water | 14.75 |
| Final volume | 25 |

| PCR program | | |
|---|---|---|
| 94 ℃ | 1 min | 1 cycle |
| 98 ℃ | 10 sec | 30 cycles |
| 68 ℃ | 15 min | |
| 72 ℃ | 10 min | 1 cycle |

PCR amplification with Herculase II

| Reaction mix | |
|---|---|
| Herculase II fusion pol | 0.5 |
| 5x Herculase II reaction buffer | 5 |
| dNTPs (10 mM each) | 0.625 |
| FW primer (2 µM) | 1.25 |
| RV primer (2 µM) | 1.25 |
| Template (20 ng gDNA) | 1 |
| Water | 15.38 |
| Final volume | 25 |

| PCR program | | |
|---|---|---|
| 94 ℃ | 2 min | 1 cycle |
| 98 ℃ | 20 sec | 30 cycles |
| $T_m$ - 5℃ | 20 sec | |
| 68 ℃ | 30 sec/1 kb | |
| 68 ℃ | 8 min | 1 cycle |

### Supplementary Note 1: Synthetic clusters used in this study

Cluster C encodes 9 genes involved in the cell cycle:

| Abbreviation | Gene |
|---|---|
| C1 | CLN2 |
| C2 | CLB2 |
| C3 | FKH2 |
| C4 | CLB3 |
| C5 | CLB5 |
| C6 | SIC1 |
| C7 | CLB4 |
| C8 | CLN1 |
| C9 | FKH1 |

Cluster H encodes 7 genes involved in biosynthesis of histidine:

| Abbreviation | Gene |
|---|---|
| H1 | HIS1 |
| H2 | HIS2 |
| H3 | HIS3 |
| H4 | HIS4 |
| H5 | HIS5 |
| H6 | HIS6 |
| H7 | HIS7 |

### Supplementary Table 1: List of primers used in this study

| Name | Sequence | Purpose |
|---|---|---|
| KC001 | CTAAACTCCATTGCTAAGGGCACTAC | Amplification of cluster integrated into YEL022W locus (primer set 1, FW) |
| KC002 | AGACACTAACACTGAGATCCCGGT | Amplification of cluster integrated into YEL022W locus (primer set 1, RV) |
| KC003 | GCTGCACAATCAACAATGATAGCCG | Amplification of cluster integrated into YEL022W locus (primer set 2, FW) |
| KC004 | ATTGTCCAACTAGCGGATACGATGG | Amplification of cluster integrated into YEL022W locus (primer set 2, RV) |
| KC005 | GTGGCTGTGGTTTCAGGGTCCA | Amplification of cluster integrated into YEL022W locus (primer set 3, FW) |
| KC006 | GAAATCATTACGACCGAGATTCCCG | Amplification of cluster integrated into YEL022W locus (primer set 3, RV) |
| KC007 | atgagtgtcagcgagtgtaactcgaGT GGCTGTGGTTTCAGGGTCCA | Amplification of histidine cluster and addition of FW barcode 1 |
| KC008 | ttgtactaatcggcttcaacgtgccGT GGCTGTGGTTTCAGGGTCCA | Amplification of histidine cluster and addition of FW barcode 2 |
| KC009 | cgcctgctacaacaggagtatcaaaGT GGCTGTGGTTTCAGGGTCCA | Amplification of histidine cluster and addition of FW barcode 3 |
| KC010 | cagaaatccgcgtgattacgagtcgGT GGCTGTGGTTTCAGGGTCCA | Amplification of histidine cluster and addition of FW barcode 4 |
| KC011 | ccctgcggtcacgtctatagaaattGT GGCTGTGGTTTCAGGGTCCA | Amplification of histidine cluster and addition of FW barcode 5 |
| KC012 | taatagcacacacggggcaataccaGT GGCTGTGGTTTCAGGGTCCA | Amplification of histidine cluster and addition of FW barcode 6 |
| KC013 | ccacagcgaggaagtaaactgttatGT GGCTGTGGTTTCAGGGTCCA | Amplification of histidine cluster and addition of FW barcode 7 |
| KC014 | cagtttaccgcatcttgacctaactGT GGCTGTGGTTTCAGGGTCCA | Amplification of histidine cluster and addition of FW barcode 8 |
| KC015 | catccaacactctacgccctcttcaGA AATCATTACGACCGAGATTCCCG | Amplification of histidine cluster and addition of RV barcode 1 |

**Supplementary Table 2: Accession numbers of raw sequencing data submitted to SRA**

BioProject PRJNA1120205

| Accession number | Sequencing description |
|---|---|
| SRR29288195 | Pool with combinatorial deletions in C cluster sequenced on Flongle |
| SRR29288194 | Pool with combinatorial deletions in C cluster sequenced on MinION |
| SRR29288193 | Post-SCRaMbLE pool with H cluster sequenced on Flongle |
| SRR29288192 | Strain1 with post-SCRaMbLE H cluster sequenced on Flongle |
| SRR29288191 | Strain2 with post-SCRaMbLE H cluster sequenced on Flongle |
| SRR29288190 | Strain3 with post-SCRaMbLE H cluster sequenced on Flongle |
| SRR29288189 | Strain4 with post-SCRaMbLE H cluster sequenced on Flongle |
| SRR29288188 | Strain5 with post-SCRaMbLE H cluster sequenced on Flongle |
| SRR29288187 | Strain6 with post-SCRaMbLE H cluster sequenced on Flongle |
| SRR29288186 | Strain7 with post-SCRaMbLE H cluster sequenced on Flongle |
| SRR29288185 | Strain8 with post-SCRaMbLE H cluster sequenced on Flongle |

**Supplementary Table 3: Median Q-scores of the data generated from described sequencing runs.**

| Sample | C cluster | C cluster | H |
|---|---|---|---|
| Flow cell | Flongle | MinION | Flongle |
| Q-score | 12.1 | | 12.5 |

Pass reads were selected based on the Q-score=9 which corresponds to 12.6% of erroneous bases in the sequence. This value was used as a threshold for annotation coverage and sequence identity. As shown in the table above, the median Q-score of the generated sequencing data and basecalled in high accuracy model exceeded 12 (error <6.3%).

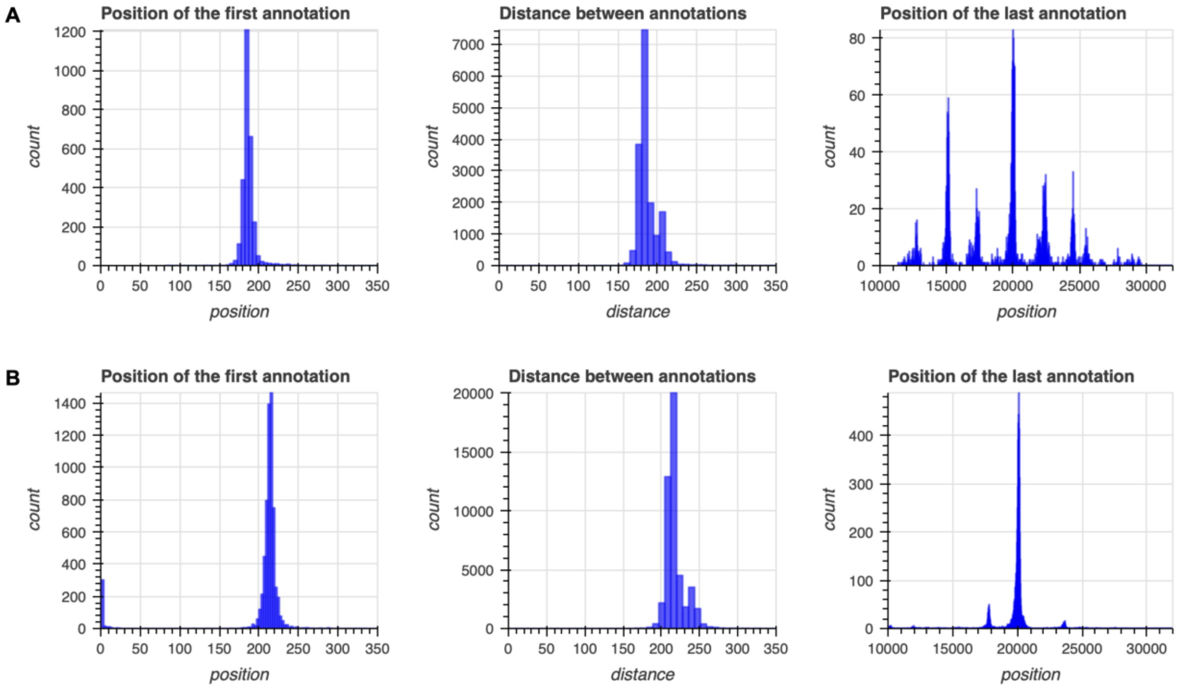


**Supplementary Figure 1: Position of annotations in the analysed reads from Flongle sequencing.** Lifted annotations include transcriptional units of cluster's genes (including promoter, gene and terminator) and the transcriptional unit of the marker. **A. Cyclins cluster**. Based on the design, the first annotation of a gene is expected at 210 bp, and the distance between annotations – 185 bp. Peaks in the plot with the position of the last annotation indicated varied size of the post-SCRaMbLEd cluster and indicated varied number of knocked-out genes. **B. Histidine cluster**. The first annotation of a gene – 239 bp, and the distance between annotations – 212-240 bp (depending on the connector). The figure with the position of the last annotation indicates that post-SCRaMbLEd clusters are approx. 17.8 kb, 20 kb or 23.5 kb.

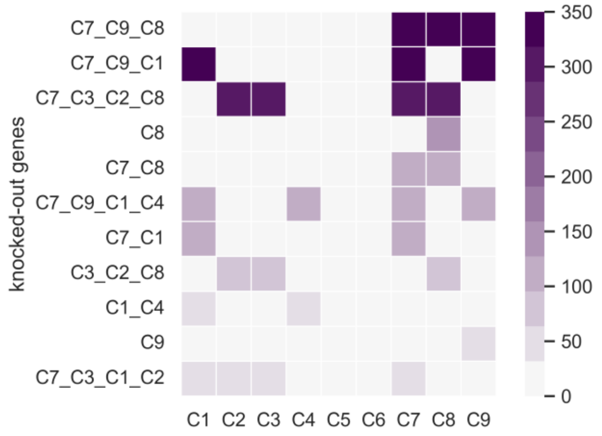


**Supplementary Figure 2: Combination of the top 11 gene knockouts in the C cluster sequenced on Flongle flow cell.**
